# Distinct afferent signatures onto anterior and posterior VMHvl subdomains and their progesterone receptor-expressing neurons

**DOI:** 10.64898/2026.05.27.728145

**Authors:** Pablo Ruiz-Campillos, Julija Raudyte, Diogo F.M. Matias, Pablo Drees-Gimenez, Inês C. Dias, Liliana Ferreira, Susana Q. Lima, Nicolas Gutierrez-Castellanos

## Abstract

Female sexual behavior is fundamentally coupled to reproductive state, requiring the dynamic and coordinated regulation of sexual receptivity and rejection across the estrous cycle. In mice, these opposing behaviors are controlled by distinct neuronal subpopulations within the anteroposterior subdomains of the ventrolateral ventromedial hypothalamus (VMHvl): the posterior VMHvl (pVMHvl) promotes sexual receptivity, whereas the anterior VMHvl (aVMHvl) drives sexual rejection. Although progesterone receptor-expressing (PR⁺) neurons in both subregions are essential for these behaviors, the circuit mechanisms underlying this anteroposterior functional segregation remain poorly understood. Here, we investigated whether anterior and posterior VMHvl subdomains receive distinct afferent inputs that could support their divergent roles, and whether these inputs specifically target PR⁺ neurons. By combining retrograde tracing with whole-brain, machine learning-assisted quantification, we first mapped the afferent connectivity of the aVMHvl and pVMHvl. We found that while the two subdomains share the majority of their inputs, they also receive distinct and biased projections. The aVMHvl preferentially receives inputs from the anterior hypothalamic nucleus, paraventricular and peripeduncular thalamic nuclei, and the parabrachial nucleus, while the pVMHvl is preferentially innervated by the preoptic area, dorsal and ventral premammillary nuclei and anteroventral periventricular nucleus. To determine whether the identified afferents directly synapse onto the PR⁺ neurons, we performed monosynaptic rabies tracing from anterior and posterior VMHvl PR⁺ neurons. These experiments revealed that most anterior- and posterior-biased projections directly targeted PR⁺ VMHvl neurons. In addition, projections from the paraventricular thalamus, periaqueductal gray, and several pontine nuclei preferentially innervated aVMHvl^PR⁺^ neurons, whereas inputs from the mammillary nuclei mostly targeted pVMHvl^PR⁺^ neurons.

Together, our results reveal distinct afferent architectures onto anterior and posterior PR⁺ VMHvl neurons, providing a circuit-level framework for understanding how hormonal state flexibly biases female sexual behavior toward receptivity or rejection.

## Introduction

Sex is pervasive in the animal kingdom; however, in most animal species, including mice, sexual behavior is not continuously expressed. Given its importance for species survival and evolutionary success, sexual behavior is precisely timed to reproductive capacity to ensure successful sexual reproduction^1–3^. In female mice, the estrous cycle gates the outcome of sexual interactions: during the fertile phase, females are sexually receptive and allow male mount attempts to progress to copulation, whereas outside this window they actively reject sexual advances, effectively preventing intercourse^1,4^. Thus, the estrous cycle coordinates a dynamic balance between sexual receptivity and competing behavioral states, such as defensive or avoidance behaviors, to flexibly allow mating only when reproductive conditions are favorable. This tight coupling between physiological fertility and behavioral receptivity is orchestrated by fluctuating levels of the female sex hormones estrogen and progesterone, which act on defined neural circuits expressing their cognate receptors^5–8^.

It is well established that the ventrolateral part of the ventromedial hypothalamus (VMHvl) is fundamental for the expression of female sexual receptivity^9,10^, a role that is specifically carried out by the posterior subregion of the VMHvl (pVMHvl)^6,7,11,12^. In contrast, its anterior counterpart (anterior VMHvl or aVMHvl) has recently been shown to control the expression of sexual rejection across the estrous cycle^4,13^. While it has been shown that progesterone receptor-expressing neurons in the anterior and posterior subdomains of the VMHvl are necessary for the expression of sexual rejection and receptivity, respectively^4,7,11^, the circuit mechanisms supporting this functional segregation along the VMHvl’s anterior-posterior (AP) axis remain poorly understood.

One possible candidate for such functional segregation is differences at the level of anatomical connections. Because functional distinctions within the VMHvl have only recently begun to receive attention, including roles in promoting mating and aggressive behaviors, as well as in mediating defensive responses to unwanted mating and aggression, as well as transcriptomic and electrophysiological heterogeneity along its AP axis^13–16^, most classical anatomical studies have not systematically examined potential AP differences in VMHvl connectivity. However, two elegant studies mapped the connectivity of estrogen receptor-expressing neurons (ER+) within the VMHvl in mice and reported pronounced AP differences in their efferent projections^14,17^. Specifically, aVMHvl^ER+^ neurons projected strongly to regions such as the periaqueductal gray, whereas pVMHvl^ER+^ neurons preferentially engaged intrahypothalamic targets, a pattern that has been recently corroborated for PR+ neurons^18^. However, whether comparable AP differences exist at the level of afferent inputs—and in particular, inputs to progesterone receptor-expressing (PR⁺) VMHvl neurons—remains largely unexplored.

Here, using a combination of retrograde tracing approaches, including retrograde transynaptic tracing^19^ from PR+ neurons, we investigated the anatomical inputs to the anterior and posterior subregions of the VMHvl. Our results show that the aVMHvl and pVMHvl receive shared inputs from ∼60% of projecting areas. However, projections from the anterior hypothalamic nucleus (AHN) and peripeduncular nucleus of the paralaminar thalamus (PP) are stronger to the aVMHvl, while projections from the preoptic hypothalamus (POA) and premammillary ventral nucleus of the hypothalamus (PMV) show a bias towards the pVMHvl. Moreover, projections from the anterior paraventricular thalamus (PVA) and parabrachial nucleus (PBN), are restricted to aVMHvl neurons and projections from anteroventral periventricular hypothalamus (AVPV) and premammillary dorsal hypothalamus (PMD) are restricted to pVMHvl neurons. Importantly, retrograde monosynaptic tracing from the anterior and posterior PR+ populations largely corroborated these different input signatures to the VMHvl AP subdomains, which could support their functional divergence in the control of female sexual behavior.

## Results

### The classic retrograde tracer Fluorogold revealed shared and exclusive inputs to the aVMHvl and pVMHvl

To map the anteroposterior afferent connectivity of the VMHvl, we performed iontophoretic injections of the classic tracer fluorogold (FG) restricted to the anterior and posterior subdomains of the VMHvl (Fig. 1A). We then employed a previously published machine learning-assisted analysis pipeline^20,21^ to perform a brain-wide quantification of FG-positive somas (FG⁺), reporting both the fraction of total input, as well as regional densities (Fig. 1B-C, Fig. 2A-J, Fig. S1 and Table 1).

**Figure 1.**
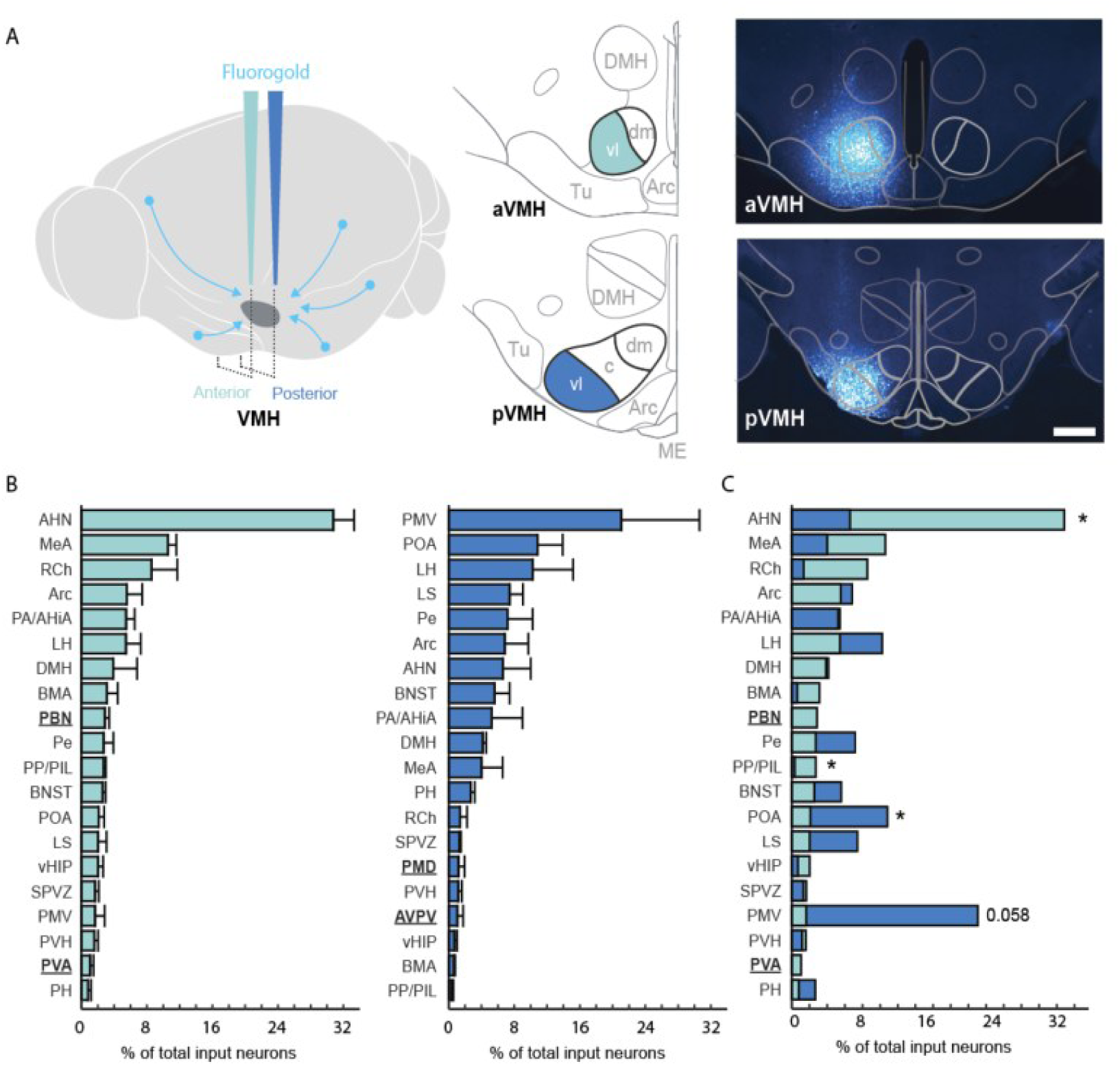
Retrograde tracing from anterior and posterior VMHvl subregions using FG, with whole-brain quantification of afferent inputs expressed as fraction of total input. (A) Schematic of the FG injection strategy (left); detailed Paxinos atlas sections showing aVMHvl and pVMHvl at anteroposterior Bregma coordinates −1.2 mm and −1.8 mm (middle); and representative FG injections in the aVMHvl and pVMHvl (right). (B) Whole-brain quantification of FG⁺ neurons following injections into the aVMHvl (left, N= 4 mice) and pVMHvl (right, N = 3 mice), performed using a machine learning-assisted analysis pipeline. Bold underlined brain areas indicate projections exclusive to the aVMHvl (left) or pVMHvl (right). The bar graph represents the mean and standard error of the mean (SEM). Overlay of both projection patterns for direct comparison is shown (C). Scale bar 400 µm. * represent p<0.05 using a permutation test.

**Figure 2.**
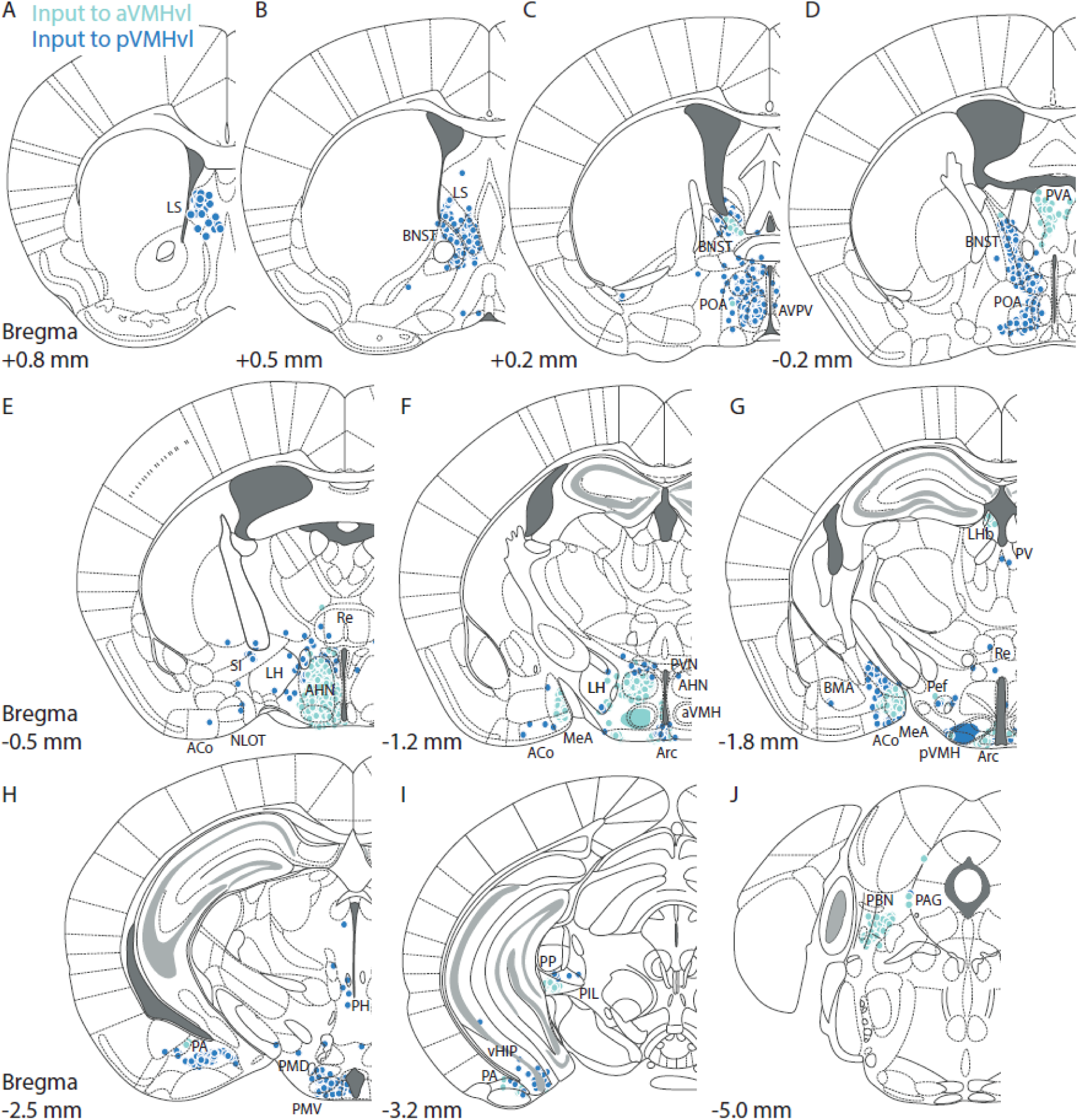
Distribution of retrograde labeling following FG injection into the aVMHvl or pVMHvl, obtained using a machine learning-based analysis pipeline. Anteroposterior sections span from the most anterior to the most posterior Bregma levels containing FG⁺ neurons. Panels show major afferent inputs from the LS (A-B), BNST and POA (C-D), AHN (E), MeA (F-G), PA and PMV (H), vHIP and PP (I), and PBN (J).

**Table 1.** Retrograde tracing from anterior and posterior VMHvl subregions using FG, with whole-brain quantification of afferent inputs expressed as projection density.

| -Region | Fraction of total input |  |  |  | Input Density (FG+ cells per um2) |  |  |  |
| --- | --- | --- | --- | --- | --- | --- | --- | --- |
| | Mean ± SEM | | p-value | z-score | Mean ± SEM | | p-value | z-score |
|  | aVMHvl | pVMHvl |  |  | aVMHvl | pVMHvl |  |  |
| **AHN** | 31.98 ± 2.52 | 6.58 ± 3.40 | **0.0302** | -2.302 | 339.95 ± 39.99 | 91.36 ± 64.38 | **0.0302** | -2.05 |
| MeA | 10.59 ± 0.98 | 4.00 ± 2.56 | 0.0603 | -1.88 | 51.09 ± 13.89 | 18.56 ± 10.52 | 0.1461 | -1.50 |
| RCh | 8.57 ± 3.13 | 1.36 ± 0.85 | 0.1159 | -1.59 | 330.44 ± 87.34 | 111.18 ± 85.27 | 0.1717 | -1.50 |
| Arc | 5.54 ± 1.89 | 6.84 ± 2.86 | 0.649 | 0.42 | 194.98 ± 59.68 | 174.75 ± 25.70 | 0.8547 | -0.29 |
| PA | 5.46 ± 1.04 | 5.24 ± 3.76 | 1 | -0.07 | 10.92 ± 4.62 | 8.24 ± 6.05 | 0.6267 | -0.38 |
| LH | 5.45 ± 1.805 | 10.24 ± 4.92 | 0.373 | 1.02 | 22.89 ± 11.70 | 42.21 ± 29.54 | 0.5761 | 0.71 |
| DMH | 3.87 ± 2.91 | 4.15 ± 0.38 | 1 | 0.08 | 100.79 ± 69.96 | 91.15 ± 20.01 | 0.9708 | -0.12 |
| BMA | 3.13 ± 1.31 | 0.62 ± 0.14 | 0.135 | -1.42 | 20.57 ± 9.38 | 3.44 ± 1.01 | 0.2173 | -1.38 |
| Pe | 2.72 ± 1.18 | 7.17 ± 3.05 | 0.2551 | 1.38 | 40.88 ± 12.78 | 93.54 ± 42.21 | 0.1741 | 1.28 |
| **PP/PIL** | 2.72 ± 0.26 | 0.35 ± 0.23 | **0.0302** | -2.31 | 87.50 ± 16.15 | 10.49 ± 6.87 | **0.0302** | -2.11 |
| BNST | 2.57 ± 0.39 | 5.59 ± 1.83 | 0.18 | 1.57 | 20.80 ± 8.97 | 41.64 ± 20.89 | 0.3798 | 1.01 |
| **POA** | 2.10 ± 0.66 | 10.82 ± 3.09 | **0.0302** | 2.01 | 17.84 ± 4.98 | 108.91 ± 60.90 | **0.0302** | 1.52 |
| LS | 2.03 ± 1.02 | 7.44 ± 1.61 | 0.0905 | 1.95 | 7.97 ± 4.68 | 23.71 ± 9.89 | 0.1784 | 1.41 |
| vHip | 2.03 ± 0.62 | 0.69 ± 0.32 | 0.1381 | -1.47 | 1.15 ± 0.49 | 0.34 ± 0.16 | 0.2211 | -1.26 |
| SPVZ | 1.65 ± 0.48 | 1.31 ± 0.16 | 0.59 | -0.62 | 105.88 ± 31.54 | 85.82 ± 25.03 | 0.713 | -0.50 |
| **PMV** | 1.65 ± 1.20 | 21.05 ± 9.71 | 0.0581 | 1.77 | 46.43 ± 24.21 | 644.39 ± 214.01 | **0.0302** | 2.02 |
| PVH | 1.59 ± 0.44 | 1.15 ± 0.46 | 0.511 | -0.70 | 42.039 ± 15.26 | 37.13 ± 24.07 | 0.8557 | -0.19 |
| PH | 0.80 ± 0.36 | 2.66 ± 0.49 | 0.0926 | 1.982 | 16.04 ± 10.35 | 29.80 ± 7.81 | 0.4038 | 0.99 |
| SCh | 0.59 ± 0.29 | 0.17 ± 0.01 | 0.2636 | -1.17 | 68.89 ± 28.11 | 17.50 ± 5.30 | 0.1455 | -1.38 |
| PPN | 0.39 ± 0.21 | .... | .... | .... | 4.02 ± 2.37 | .... | .... | .... |
| PBN | 2.87 ± 0.53 | .... | .... | .... | 24.02 ± 6.62 | .... | .... | .... |
| PVA | 1.06 ± 0.42 | .... | .... | .... | 15.97 ± 4.98 | .... | .... | .... |
| PT | 0.27 ± 0.09 | .... | .... | .... | 8.73 ± 3.66 | .... | .... | .... |
| Re | 0.25 ± 0.09 | .... | .... | .... | 5.34 ± 2.89 | .... | .... | .... |
| AVPV | .... | 1.05 ± 0.70 | .... | .... | .... | 63.89 ± 53.78 | .... | .... |
| PMD | .... | 1.19 ± 0.73 | .... | .... | .... | 68.25 ± 37.52 | .... | .... |
| TT | .... | 0.23 ± 0.06 | .... | .... | .... | 1.19 ± 0.03 | .... | .... |

Our results reveal a subset of projections that exclusively target either the aVMHvl or the pVMHvl, a subset that targets both regions but exhibits a significant AP bias, and a large subset that targets both AP levels comparably (Fig. 1B-C). Two thalamic projections, arising from the paralaminar thalamus and the anterior paraventricular nucleus of the thalamus (PVA), selectively targeted anterior levels of the VMHvl, with little to no innervation of posterior regions (Fig. 1B, Fig. 2D,I). Within the paralaminar thalamus, projections from the peripeduncular thalamus (PP) were particularly dense and innervated exclusively the aVMHvl. In addition, a small number of FG⁺ neurons were detected in the adjacent posterior intralaminar thalamic nucleus (PIL) (Fig. 2I). In contrast, injections confined to the pVMHvl labeled only sparse PIL neurons, resulting in a significant AP bias (permutation test, Observed Standardized Statistic Z=-2.11, permutations=10,000, p=0.03). In addition to these thalamic inputs, a projection arising from the lateral portion of the parabrachial nucleus (PBN) also specifically targeted anterior, but not posterior, VMHvl levels (Fig. 1B, Fig. 2J).

In contrast, the projections from the anteroventral paraventricular hypothalamus (AVPV) and dorsal premammillary nucleus (PMD) showed a strong preference for innervating the pVMHvl, mostly sparing the anterior domain (Fig. 1B, Fig. 2B-C, Fig. 2G-H).

Three intrahypothalamic projections showed a significant AP bias (Fig. 1C). The projection from the anterior hypothalamic nucleus (AHN, permutation test, Z=-2.09, permutations=10,000, p=0.03) was significantly denser onto the aVMHvl (Fig. 1C, Fig. 2E). In contrast, the preoptic area of the hypothalamus (POA, Z=1.52, permutations=10,000, p=0.03) showed a significant pVMHvl preference (Fig. 1C, Fig. 2C, D, H). Finally, the ventral premammillary nucleus showed a marginally non-significant bias towards the pVMHvl (PMV, Z=1.77, permutations=10,000, p=0.05); however, this bias reached significance when projection density was compared (PMV, Z=2.02, permutations=10,000, p=0.03).

Several regions showed trends toward an AP bias. Specifically, the medial amygdala (MeA), retrochiasmatic area (RCh), and basomedial amygdala (BMA) exhibited a visible projection bias toward the aVMHvl, whereas the bed nucleus of the stria terminalis (BNST), lateral septum (LS), and the lateral, posterior, and periventricular hypothalamic nuclei (LH, PH, and Pe, respectively) showed a visible preference for projecting to the pVMHvl. However, none of these differences reached statistical significance with the current sample size (Fig. 1B-C, Table 1).

Finally, several intrahypothalamic projections, including those from the arcuate nucleus (Arc), subparaventricular zone (SPVZ), dorsomedial hypothalamus (DMH), and paraventricular hypothalamus (PVH), as well as extra-hypothalamic inputs from the posterior amygdala/amygdalohippocampal transition area (PA/AHiA) and ventral hippocampus (vHIP), showed no AP bias and projected in a comparable manner to both anterior and posterior VMHvl populations (Fig. 1B-C, Table 1).

Given that our analysis pipeline considered only brain nuclei and did not resolve subnuclear differences (e.g., dorsal vs. ventral medial amygdala), we next examined the spatial distribution of all identified projections along the anteroposterior, mediolateral, and dorsoventral axes in representative aVMHvl and pVMHvl injections (Fig. 2), which revealed pronounced spatial biases within several regions, including the BNST, POA, and MeA. We therefore performed a more detailed analysis based on manual scoring of FG⁺ labeled neurons to further characterize these projections at the level of their constituent subdivisions.

Our results revealed robust differences in the pattern of aVMHvl and pVMHvl innervation across spatially segregated subdivisions of the MeA, POA, and BNST (Fig. 3). Within the MeA, the anterodorsal and posteroventral subdivisions preferentially innervated the aVMHvl, whereas all other MeA subdivisions examined showed no AP bias (Fig. 3A). In the preoptic hypothalamus (POA, scored at Bregma level -0.1mm), projections from the periventricular hypothalamic nucleus (Pe) and the medial and lateral parts of the medial preoptic nucleus (MPOM and MPOL, respectively) were significantly denser onto the pVMHvl compared to the aVMHvl. In contrast, projections from the medial preoptic area (MPA) and the ventrolateral and lateral subdivisions of the preoptic nucleus (VLPO and LPO, respectively) showed no AP bias (Fig. 3B). In the BNST, only the postero-medial subdivision produced a significantly denser projection to the pVMHvl than to the aVMHvl, while all other subdivisions showed no AP bias (Fig. 3C).

**Figure 3.**
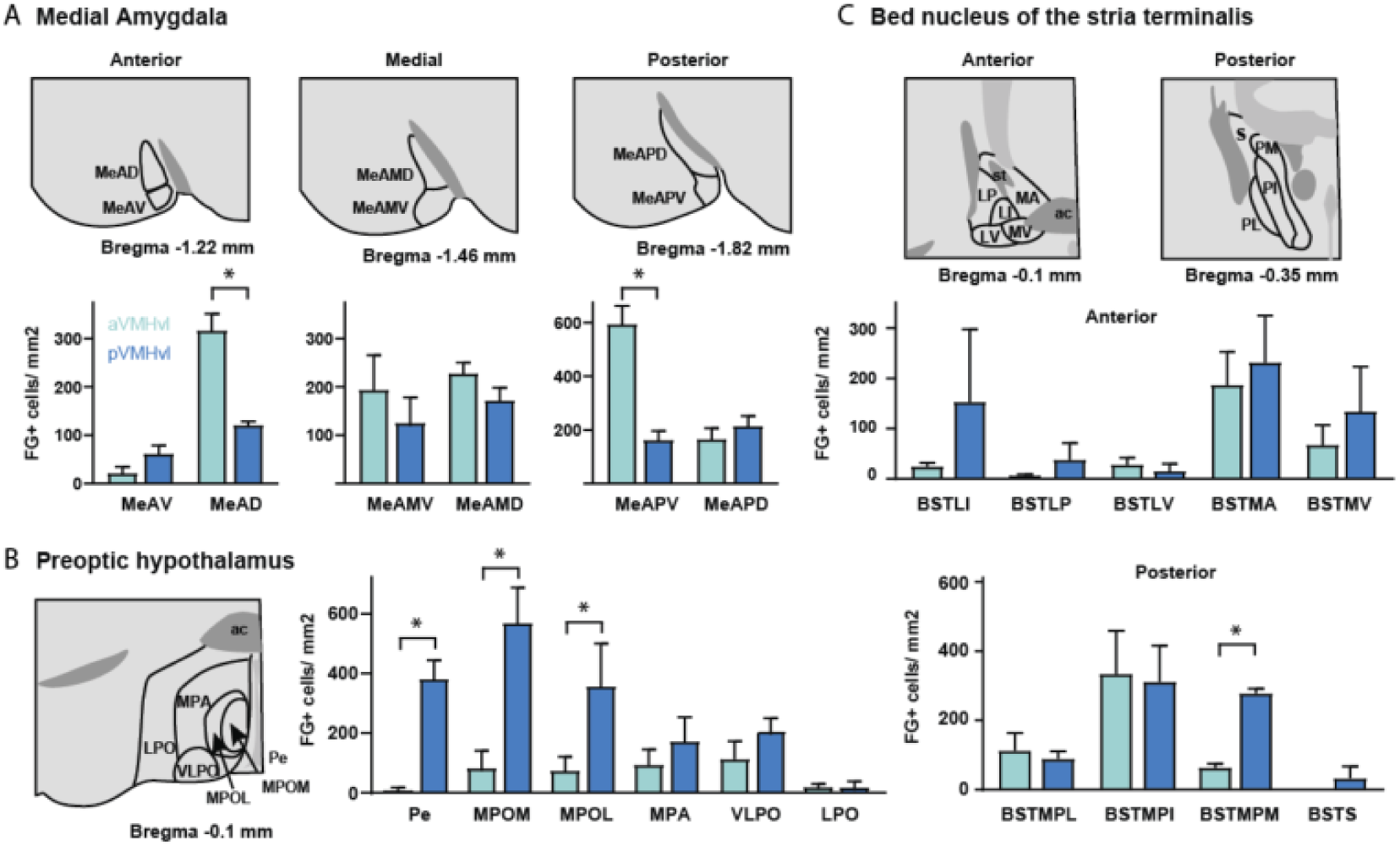
Subnuclear organization of afferent inputs to aVMHvl and pVMHvl. (A) Schematic representation of the different subdivisions considered for the medial amygdaloid nucleus at its different anteroposterior levels (top). Quantification of the density of FG⁺ cells in each subdivision for aVMHvl and pVMHvl FG injections. (B) Same as in (A), shown for the Preoptic hypothalamus. (C) Same as in (A), shown for the Bed nucleus of the stria terminalis. * represent p<0.05 using Sidak’s test.

### Rabies assisted transynaptic retrograde tracing revealed shared and exclusive input patterns to PR+ neurons in the aVMHvl and pVMHvl

Classic retrograde tracers such as FG present several caveats. For example, these tracers do not specifically label projections to a genetically defined neuronal subpopulation, as they unspecifically label all afferent projections to the injection site. Importantly, they can also label axons of passage: fibers that do not project to the injection area but simply pass through it. This may be worth considering in the VMH, which is embedded within a dense terminal field containing axons from the stria terminalis, mammillary peduncle, medial forebrain bundle, lateral hypothalamus, and fibers entering the diencephalon through the zona incerta^22^, some of which may not form synaptic contacts with VMH neurons. To circumvent these issues, we sought to trace the monosynaptic afferent inputs to the PR⁺ population of the VMHvl using the Avian Sarcoma Leukosis Virus Subtype A (EnvA) G-deleted rabies virus system (ΔG-RV)^19^.

To achieve this, we first delivered a combination of Cre-dependent adenoassociated viruses (AAVs) carrying the avian viral receptor TVA, the reporter mCherry, and the viral glycoprotein G into female mice expressing Cre under the progesterone receptor promoter (PR-Cre^11^). This was followed by the delivery of ΔG-RV particles expressing GFP to the same location (Fig. 4A; see Methods). With this approach, mCherry expression defines the population of cells that can be infected by the rabies virus via the TVA receptor. In this population, expression of the glycoprotein G enables the assembly of new viral particles that travel retrogradely to presynaptic neurons. This approach results in three distinct fluorescently-labeled populations: red-only cells, which express mCherry but were not infected by the rabies virus; yellow cells, which co-express mCherry and GFP at the injection site and represent the starter cell population infected by rabies; and green-only cells, corresponding to retrogradely labeled presynaptic neurons arising from both local and long-range projections.

**Figure 4.**
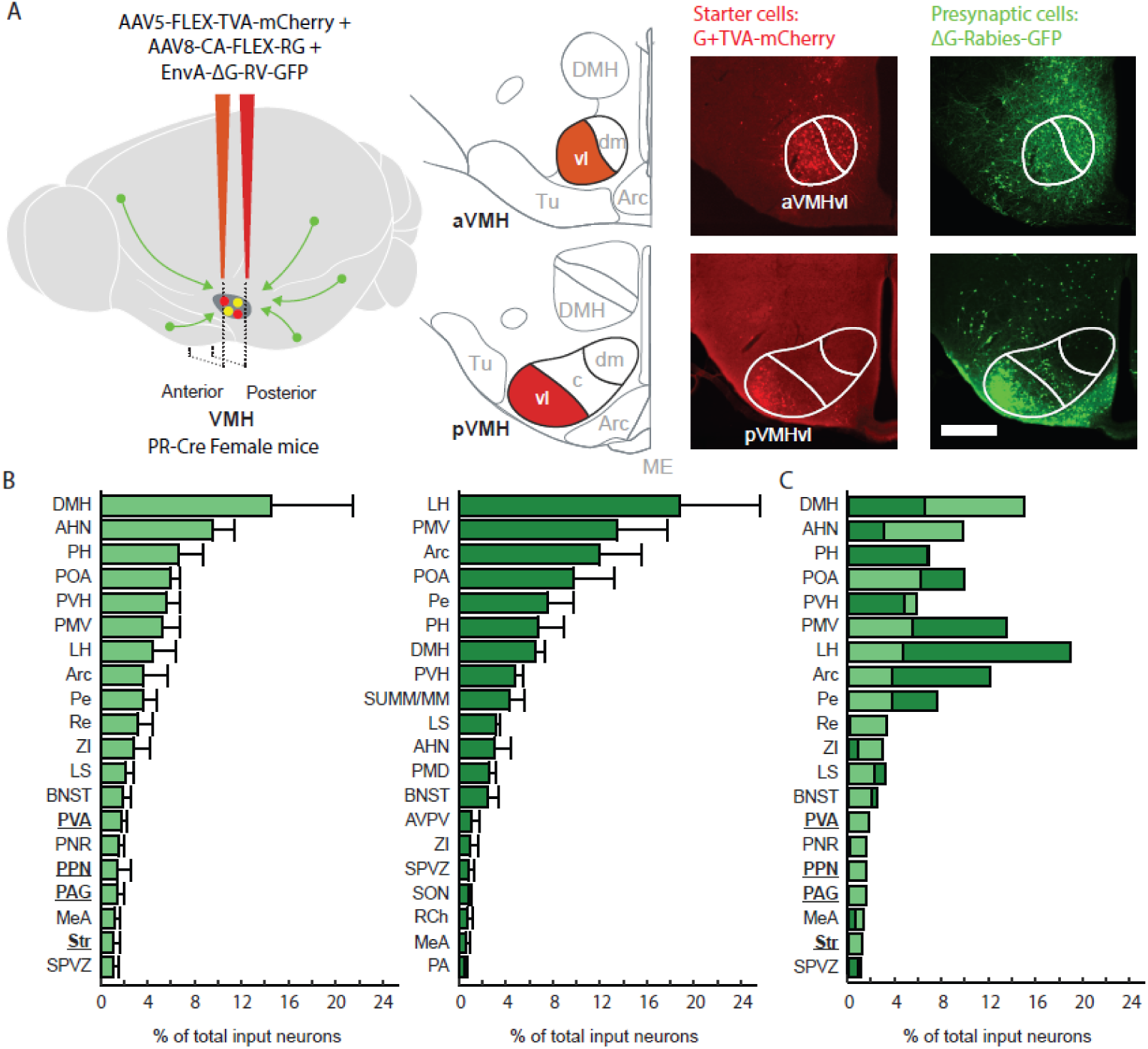
Tracing of monosynaptic inputs to anterior and posterior VMHvl PR⁺ neurons using G-deleted rabies virus. (A) Experimental design for retrograde labeling of presynaptic partners of PR⁺ neurons in the anterior or posterior VMHvl (left and middle). Representative images showing the potential starter cell population (mCherry⁺) and presynaptic neurons labeled by rabies virus (GFP⁺) (right). (B) Whole-brain quantification of rabies-labeled neurons following injections into the aVMHvl (left) and pVMHvl (middle), performed using a machine learning-assisted analysis pipeline. The bar graph represents the mean and standard error of the mean (SEM). (C) Overlay of both projection patterns for direct comparison is shown (right). Scale bar 400 µm.

After confirming that the starter population was restricted to PR⁺ neurons located either in the aVMHvl or pVMHvl (Fig. 4A), we performed a whole-brain quantification of the number of projecting neurons labeled with GFP (Fig. 4B, Table 2). Although injections in both groups were centered on the intended target regions (Fig. S2), pVMHvl injections yielded visibly larger, but not significantly different, starter cell populations compared to aVMHvl injections (Mean±SEM: aVMHvl=18±7.5, pVMHvl=67.7±39.7, Mann-Whitney, p=0.2). Because injection volumes were matched across groups, these differences likely reflect a higher density of PR⁺ neurons in the pVMHvl relative to the aVMHvl^13^. Given this, we considered comparisons based on fraction of total input to be more appropriate than projection densities, which are likely more sensitive to differences in injection size.

**Table 2.**
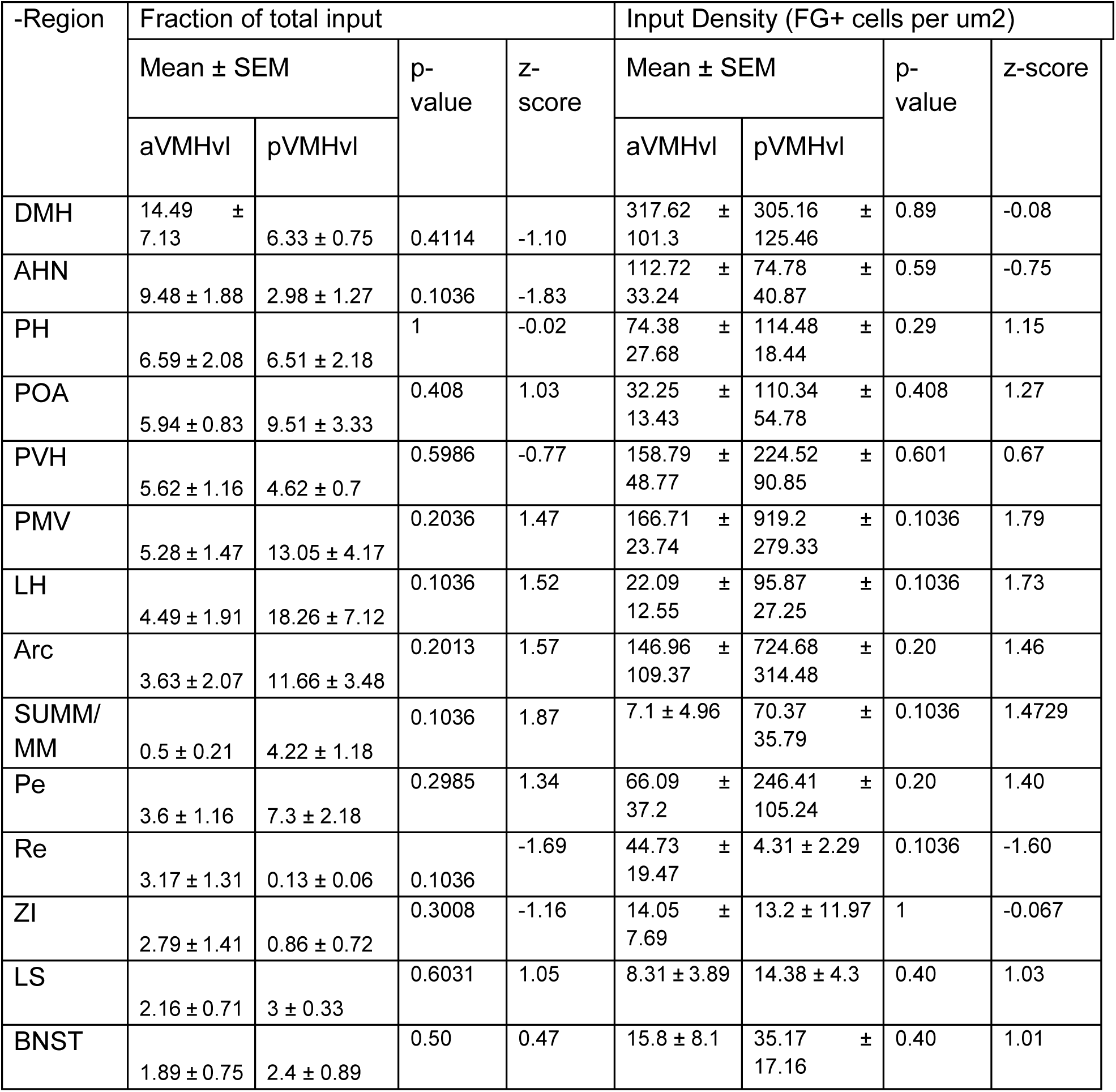
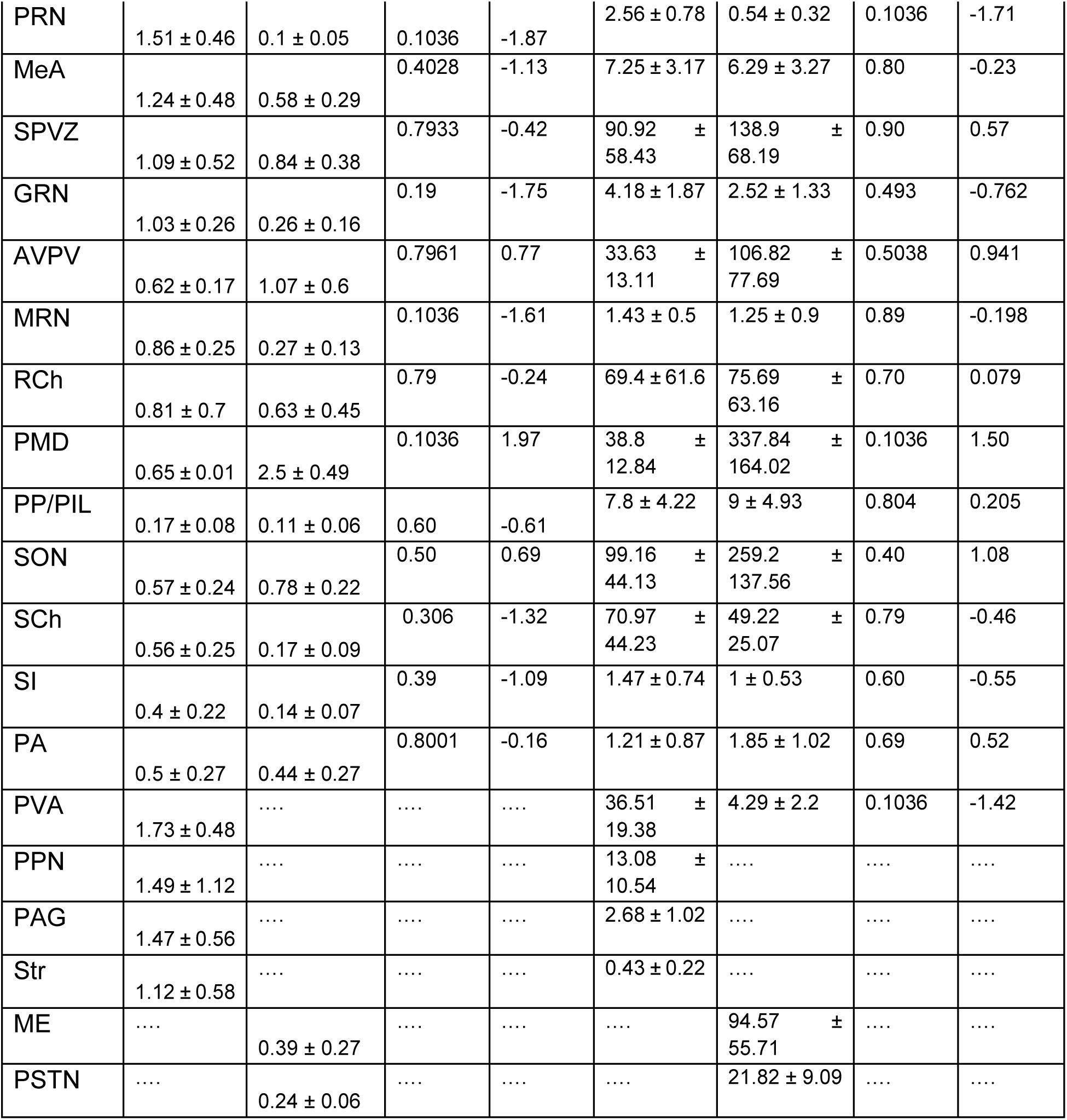
Retrograde tracing from anterior and posterior VMHvl PR+ neurons using monosynaptic retrograde tracing and whole-brain quantification of their afferent projecting areas as projection density.

Our results showed that, in line with the FG experiments, the majority of projections were shared between the two subregions (Fig. 4C and Table 2). Importantly, rabies-assisted transsynaptic retrograde tracing showed that nearly all regions identified with FG as projecting to the aVMHvl and pVMHvl, targeted at least in part PR⁺ neurons, with the BMA as the sole exception (Fig. 5). Although consistent AP bias trends were observed between both datasets—for example, preferential projections from the AHN to the aVMHvl and from the POA and PMV to the pVMHvl—these differences did not reach statistical significance (Fig. 4C, Table 2).

**Figure 5.**
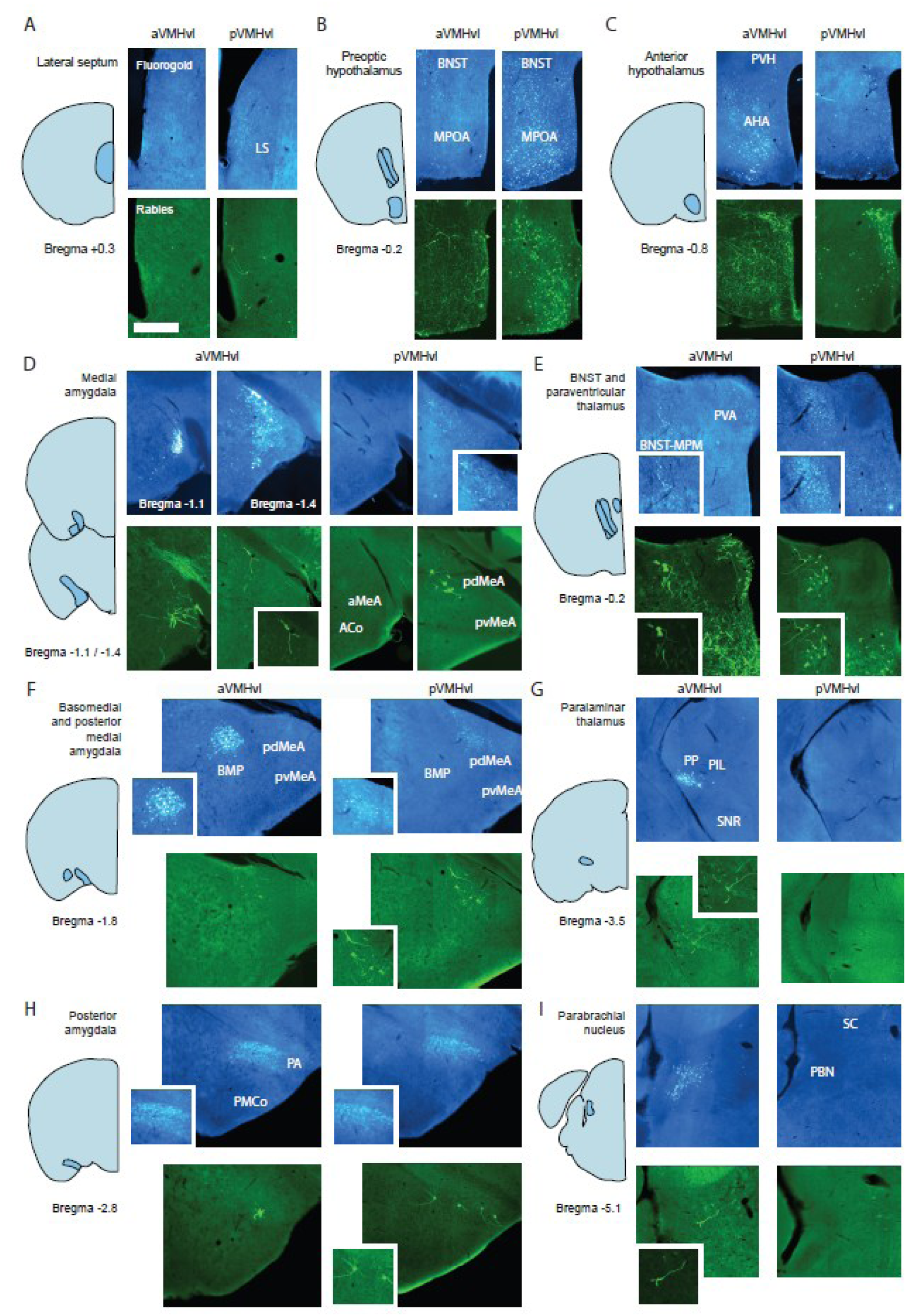
Comparison of inputs to the anterior and posterior VMHvl identified by FG and monosynaptic tracing from PR⁺ neurons. The lateral septum (A) and preoptic hypothalamus (B) project strongly to the pVMHvl, but not to the aVMHvl. In contrast, the anterior hypothalamus (C), anterior medial amygdala (D), paraventricular thalamus (E), basomedial posterior amygdala (F), and peripeduncular portion of the paralaminar thalamus (G) preferentially project to the aVMHvl. The posterior amygdala showed no clear bias in its projections (H), whereas the parabrachial nucleus preferentially projects to the aVMHvl (I). In all panels, the top images show FG tracing and the bottom images show rabies tracing. For each panel, the left plots correspond to tracing from the aVMHvl and the right plots to tracing from the pVMHvl. With the exception of the basomedial amygdala, all projections identified by FG labeling were also detected by rabies tracing. Scale bar 500 µm.

Additionally, approximately 20% of projections selectively targeted either PR⁺ neurons of the aVMHvl or pVMHvl. These included projections previously identified by FG tracing as aVMHvl-specific, such as inputs arising from the PVA. Rabies tracing further revealed additional input regions that were likely below the detection threshold of the FG analysis due to their lower projection density. These included the thalamic nucleus reuniens (Re); the striatum (Str) and the zona incerta (ZI); and several brainstem regions, including the pontine reticular nucleus (PNR), pedunculopontine nucleus (PPN), and periaqueductal gray (PAG), all of which preferentially projected to aVMHvl PR⁺ neurons. In contrast, hypothalamic regions such as supraoptic nucleus (SON), and the mammillary and supramammillary nuclei (SUMM/MM); the median eminence (ME) and parasubthalamic nucleus (PSTN) preferentially innervated pVMHvl PR⁺ neurons (Fig. 4C, Fig. 6, Fig. S3 and Table 2).

**Figure 6.**
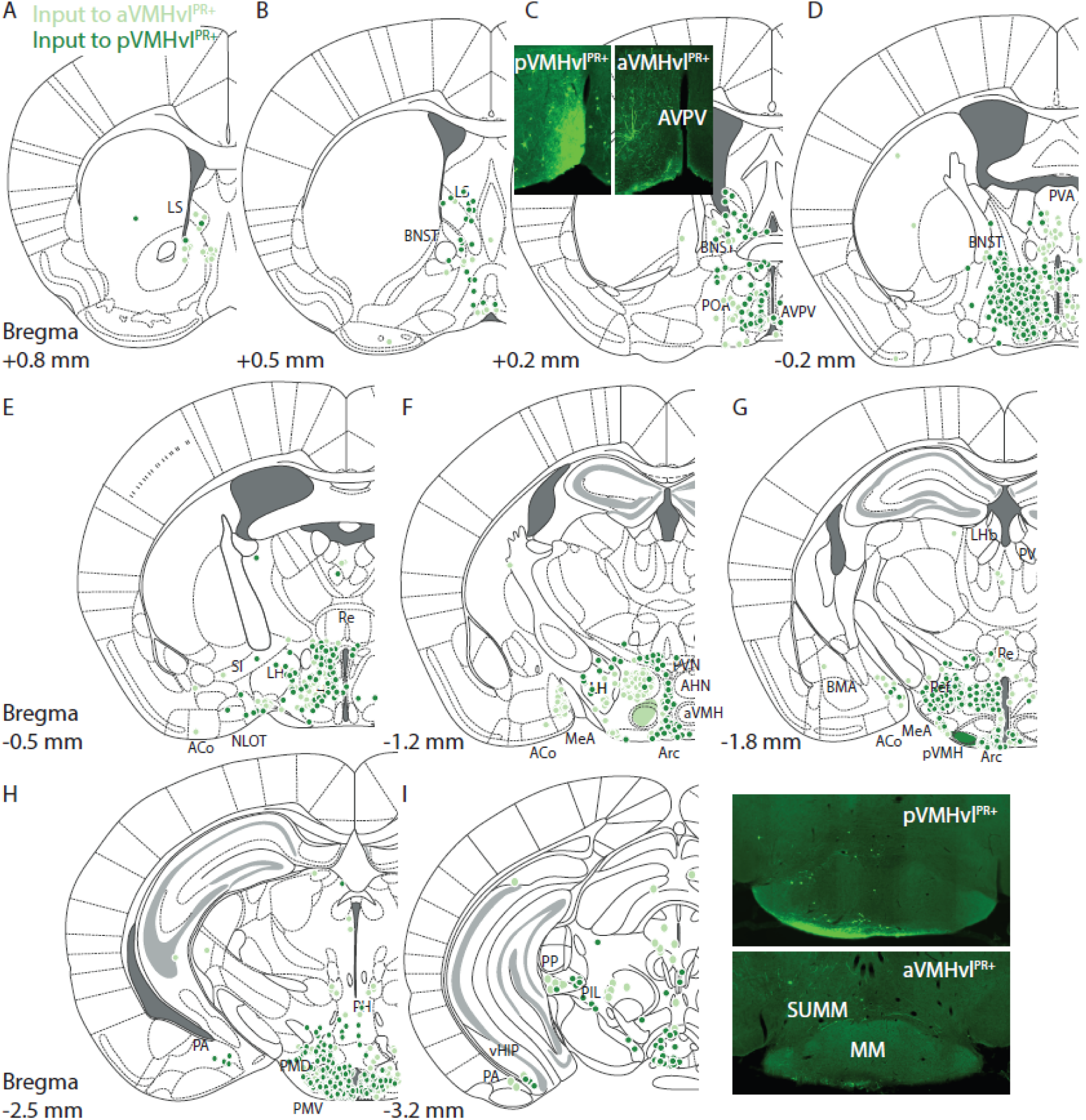
Distribution of aVMHvl and pVMHvl PR+ presynaptic partners labeled using monosynaptic rabies virus tracing. Anteroposterior sections span from the most anterior to the most posterior Bregma levels containing rabies-labeled neurons. Panels show major afferent inputs from the LS (A-B), BNST and POA (C-D), AHN and PVA (E), MeA and PVN (F-G), PA and PMV (H), and vHIP, PP and SUMM/MM (I). Quantification obtained using a machine learning based pipeline.

## Discussion

In this study, we quantitatively mapped the whole-brain afferent connectivity of the anterior and posterior subregions of the VMHvl using non-specific retrograde tracers. We then used monosynaptic retrograde viral tracing to identify which of these projections specifically synapse onto PR-expressing neurons, a population critically involved in female sexual rejection within the aVMHvl and in receptive behavior within the pVMHvl^4,7,11^.

Our results are in vast agreement with previous studies mapping the afferent connectivity pattern on the VMHvl in rodents^17,23^. However, most previous studies had not carefully taken into consideration the different AP levels of the VMHvl. To the best of our knowledge, a single study has carefully taken into consideration AP levels when examining inputs to the VMHvl^17^. In this study, an indirect intersectional retrograde tracing strategy was used to compare inputs to the anterior and posterior VMHvl by leveraging known differences in PAG- and MPOA-projecting VMHvl populations. Their results indicated largely shared presynaptic inputs to both regions, with suggestive but non-significant differences including stronger AHN input to the aVMHvl and BNST input to the pVMHvl, which are in full agreement with our observations.

Using FG, our results identified ∼60% of AP shared projections (those not yielding significant AP differences), ∼25% of anterior preferred and ∼15% of posterior preferred (projections that either showed a significant AP bias or targeted exclusively an AP subregion). Similarly, using rabies retrograde monosynaptic tracing from the PR⁺ population, we obtained ∼80% of AP shared projections, ∼15% of anterior preferred and ∼5% of posterior preferred. Importantly, “exclusive connections” refer to projections detected only for one AP level of the VMHvl, whereas “non-present connections” denote pathways that did not meet our inclusion threshold in either AP level (see methods). These categories may therefore reflect low or sparse labeling density in our dataset rather than true anatomical absence. Indeed, several such connections were detected in our FG experiments below the established cutoff, for instance inputs from the medial prefrontal cortex, some pontine nuclei, the median raphe nucleus (MRN) and the PAG. As a result, these projections could not be reliably quantified in the present analysis of global connectivity using FG, despite having been reported in previous studies as modest sources of input to the VMHvl^17^. However, most of these connections, including inputs from PAG, SUMM/MM, PRN, MRN, or SON were detected using our rabies tracing approach, suggesting that although they may not constitute major sources of input to the VMHvl as a whole, they consistently innervate its PR⁺ neuronal subpopulation.

Among the input sources that preferentially targeted the aVMHvl, we identified several nuclei known to play roles in defensive behaviors, such as the AHN, the PBN and the PVA^24^. For instance, GABAergic neurons in the AHN mediate active defensive behaviors against noxious somatosensory stimuli^25^, while the lateral PBN is linked to arousal and defensive behavior, including avoidance and escape, through calcitonin gene-related peptide- and cholecystokinin-expressing neuronal populations^23–25^. Similarly, the PVA^29^ has been shown to control the balance between active and passive defensive strategies via its GABAergic and glutamatergic subpopulations.

We also observed differential innervation from the MeA, with its anterior and posteroventral subregions preferentially innervating the aVMHvl. This is consistent with previous reports showing denser projections from the anterior MeA to the aVMHvl relative to the pVMHvl^30^. Functionally, the anterior MeA receives inputs from the main olfactory bulb and is activated by sexually relevant volatile cues^31^, whereas the posterior MeA primarily receives direct inputs from accessory olfactory bulb neurons involved in pheromone sensing^32^. Together, these findings suggest that volatile olfactory cues may exert a stronger influence over aVMHvl-dependent behaviors, whereas pheromonal information may preferentially engage the pVMHvl. Supporting this interpretation, we found significantly denser projections from the posteromedial BNST subdivision, a major recipient of accessory olfactory bulb input^33^, to the pVMHvl compared to the aVMHvl. Finally, the posteroventral MeA, which preferentially innervated the aVMHvl, has been implicated both in defensive behaviors and anxiety-like states^34,35^. However, this subnucleus of the amygdala is also engaged when in contact with the sex promoting pheromone ESP1^36^ and may convey information about conspecific individual identity^37^. Moreover, within the paralaminar thalamus, we identified a projection from the PP that preferentially targets the aVMHvl. Classic studies linked PP integrity to the expression of lordosis behavior in ovariectomized female rats primed with sex hormones^38,39^, although these findings were not replicated in naturally cycling females^40^. Future studies should investigate the role of PP inputs in female sexual behavior and other behaviors regulated by the aVMHvl.

In addition to the previously described projection from the PVA, rabies tracing revealed substantial inputs from midbrain regions, including the PAG and PPN, specifically to aVMHvl PR⁺ neurons. These premotor areas receive input from aVMHvl PR⁺ neurons^18^, likely linking activity of this subpopulation to the execution of motor outputs. In fact, optogenetic stimulation of aVMHvl PR⁺ neurons input to the dorsomedial PAG drives female sexual rejection^18^. Their reciprocal projections to the aVMHvl may reflect feedback signaling related to premotor command activity.

Among input sources that preferentially targeted the pVMHvl, we identified the preoptic hypothalamus, with significant differences observed across its medial, lateral and periventricular subdivisions. The medial preoptic area controls maternal behavior, maternal motivation and pup care through GABAergic and VGlut+ populations that can co-express markers such as galanin or estrogen receptors^41–44^. In the context of female sexual behavior, its GABAergic population has been shown to be activated upon male ejaculation and exert inhibitory effect on female sexual drive during the post-mating refractory phase^45^. The lateral preoptic area is activated during sexual behavior, although its activity is not modulated by male ejaculation^45^. Its connectivity pattern, particularly its strong inputs to VTA neurons, suggests a potential role in the motivational aspects of sexual behavior^46^.

In addition, we found that both areas composing the rostral periventricular area of the third ventricle (RP3V: AVPV and Pe) preferentially targeted the pVMHvl. The RP3V is an area relevant for controlling luteinizing hormone secretion and maintaining the estrous cycle^47^, The AVPV is the recipient of dense projections from pVMHvl PR⁺ neurons, and this pathway undergoes sex hormone-dependent structural potentiation, crucial for the expression of female receptivity^7^. The role of this feedback projection to the pVMHvl should be the focus of future research. We also observed a biased projection from the PMV, a region linked to maternal aggression^48^ via its dopamine transporter positive subpopulation (DAT+). This is consistent with its known role in males, where it controls intermale aggression, and strongly projects to the pVMHvl^49^. Given that the VMHvl contains genetically and spatially defined ensembles involved in aggression and mating^6,12,50^, it is tempting to speculate that PMV inputs may preferentially target neurons controlling maternal aggression, although this will require future investigation.

Among the projections consistently identified by both FG and rabies tracing, and representing major inputs across the AP axis of the VMHvl, we identified several regions involved in the regulation of metabolic state, hunger and satiety, and associated food seeking behavior, including the Arc, PVH, and LH ^51–54^, which also mediate behavioral selection under competing needs ^52,55^. We also observed strong inputs from the DMH, which may represent an important relay in VMH output pathways, as proposed in classic models of VMH information flow^56^. Finally, we identified inputs from the PA, a region implicated in the control of aggression and mating behavior through distinct neuronal subpopulations in male mice^57,58^. In females, elegant recent work by Yamaguchi and colleagues further demonstrated a critical role for the PA in maternal aggression, supported by transient neural adaptations that enhance activation of this pathway during the postpartum period^59^.

In most cases, rabies tracing and FG injections produced similar labeling patterns, confirming that the major projections identified with FG target, at least in part, synapsed onto PR⁺ VMHvl neurons. The main exception was the BMA, which showed dense FG labeling but no labeled neurons when the rabies strategy was employed. Since previous optogenetic assisted circuit mapping and slice electrophysiology studies demonstrated monosynaptic connections from the BMA onto VMHvl glutamatergic neurons^58^, the absence of rabies-mediated labeling suggests that BMA afferents target PR- neurons. However, given that some studies have shown undersampling of presynaptic neurons with rabies tracing^61^, the absence of rabies-mediated labeling in our dataset warrants further scrutiny.

Comparison of our input mapping with a recent characterization of PR⁺ VMHvl output projections^18^ revealed that several AP biased inputs also exhibit corresponding output biases, suggesting the presence of reciprocal functional loops. For example, the AHN, PP, and PVA, which preferentially project to the aVMHvl, are also prominent targets of aVMHvl PR⁺ neurons. Conversely, the PMV, PMD, and SUMM/MM, which preferentially innervate the pVMHvl, are major output targets of pVMHvl PR⁺ neurons.

The present study presents some technical limitations to discuss. Due to its proximity from the FG injection site, the tuberal hypothalamus exhibited strong labeling that could reflect either injection leakage or genuine retrograde transport (Fig. 1A). After carefully inspecting the injection sites of the rabies experiments, in which the starter population giving rise to the traced projections was quantitatively well defined (Fig. 4A), we observed that tuberal areas presented a modest to low density of rabies traced neurons in the total absence of starter cells. These results confirm that tuberal areas project to VMHvl PR+ neurons, but suggest either tuberal areas present stronger projections to non-PR⁺ populations within the VMHvl, or that FG injections presented a mild leakage into adjacent tuberal areas, a problem that our rabies tracing experiments did not suffer from. Also, as previously discussed, undersampling with the rabies strategy could also explain this result^61^. Because these possibilities could not be conclusively distinguished, the tuberal hypothalamus was excluded from analysis, consistent with previous studies^17^. Given that a similar concern could arise from other areas close to the injection sites, such as the arcuate, anterior or premammillary nuclei of the hypothalamus, we conducted extra analysis to rule out that injection contamination could explain some of the labeling reported in our study. For that, we carefully examined injections where no starter cells could be found in these adjacent areas (Fig. S2). Even in those cases, both the AHN and PMV still ranked among the top projecting areas to the aVMHvl and pVMHvl. These results are also consistent with studies previously reporting dense projections from the AHN^62,63^, Arc^64^, and PMV^65^ to the VMHvl. Finally, we noted a minor technical limitation of the QUINT analysis pipeline in regions with very high projection density. In these densely labeled areas, clusters of closely adjacent projection-labeled neurons were occasionally segmented as single large cells, leading to a slight tendency toward undersampling. As a result, some of the differences reported in the most densely labeled regions may be mildly underestimated.

FG tracing revealed overall denser projections than rabies-based tracing (Table 1 and 2). Although these differences may partly reflect the fact that rabies selectively traces inputs to a specific VMHvl subpopulation, they may also result from the lower labeling efficiency of the viral approach^61^. Therefore, to more precisely characterize the connectivity between the VMHvl and its afferent input regions, future studies should incorporate complementary approaches, such as optogenetically assisted circuit mapping^66^ or next-generation rabies tracing methods with improved efficiency^67^.

While we were able to identify the source of many input regions onto different domains of the VMHvl, in this study we did not investigate the molecular nature of the projecting neurons. Previous elegant work by Lo and colleagues^17^ showed that projections arising from the LS, AHN, BNST, MeApd, DMH, and ZI are predominantly GABAergic, whereas those from the PVN, AVPV, PMV, PA and hippocampus are predominantly glutamatergic. In contrast, projections from the POA and PAG showed no clear neurochemical bias^17^. However, these experiments did not take AP differences within the VMHvl into account and likely did not capture projections targeting its most rostral portion.

In the present study we show evidence for specific and poorly characterized projections from the PVA, PBN, and anterior MeA that preferentially target the aVMHvl. Future studies will be required to determine the neurochemical identity of these projection neurons and to establish their contribution to female sexual behavior through pathway specific manipulation experiments.

Altogether, our results revealed critical differences between the inputs received by the aVMHvl and pVMHvl. Recent studies have highlighted a clear functional segregation along the VMHvl AP axis, with the aVMHvl implicated in self-defense behaviors and the pVMHvl in sexual receptivity. The existence of distinct input architectures onto these subdomains composed of neuronal populations with rich expression of sex hormone receptors, provides a circuit-level framework for understanding how hormonal signals coordinate multiple aspects of female socio-sexual behavior, coupling reproductive state with the appropriate behavioral responses throughout the estrous cycle.

## Methods

### Animals

Data were collected from adult female mice (2-4 months) from the C57BL/6 strain (JAX stock #000664) for Fluorogold experiments and from B6129S(Cg)-Pgrtm1.1(Cre)Shah/AndJ75 (PR-Cre; JAX stock #017915), expressing Cre recombinase under the control of the progesterone receptor (PR) gene for monosynaptic tracing experiments.

Animals were kept under controlled temperature of 23±1C, reversed photoperiod of 12-h light/dark cycle (light available from 8 pm to 8 am) and group-housed conditions in standard cages with environmental enrichment elements. Food and water were provided ad libitum. Females were weaned at 20-21 days of age and group-housed with two to five animals. After reaching 6 weeks of age, females were exposed to adult C57BL/6 male soiled bedding once per week to stimulate the natural reproductive cycle.

Procedures were executed in accordance with the standards approved by the Commission for Experimentation and Animal Welfare of the Champalimaud Centre for the Unknown (Órgão para o Bem Estar Animal; ORBEA) and by the Portuguese National Authority for Animal Health (Direcção Geral de Alimentação e Veterinária; DGAV) (Ref. 0421/000/000/2018).

### Stereotaxic injections and histological processing

All surgeries were conducted in a stereotaxic frame (Kopf instruments) and under isoflurane anesthesia. Stereotaxic coordinates for FG injections in the aVMHvl were AP: 1.10, ML: -0.5, DV: 5.60 (N = 4) and AP: 1.5, ML: -0.75, DV: 5.60 for pVMHvl injections (N = 3). As previously described^68^, FG was injected by ionophoresis through glass micropipettes with an inner diameter tip of ∼20 µm filled with 2% FG in saline solution.

To do so, positive pulses (10s on/off; 5 µA) were applied over 3-5 min using a microstimulator (Iso-Flex, A.M.P.I). A continuous negative retaining current (0.9 µA) was applied during entrance and withdrawal of the micropipette to avoid diffusion of the tracer.

After 7 days of survival, animals received an overdose of sodium pentobarbital and were briefly perfused transcardially with PBS 0.1M, followed by 4% paraformaldehyde in 0.1 M PB (pH 7.6) at a rate of ∼5 mL/min for 10 minutes. Brains were carefully removed from the skull and post-fixed 2-3 hours in the same fixative solution at 4°C. Then, they were immersed in 30% sucrose in PB at 4°C until they sank, and 40-µm-thick coronal sections obtained with a freezing microtome were collected into four matching series.

To visualize monosynaptic inputs to VMHvl^PR+^ neurons, viral volumes were injected with a Nanoject II Auto-Nanoliter Injector (Drummond Scientific). First, a combination of AAV5-FLEX-TVA-mCherry and AAV8-CA-FLEX-RG were delivered either in the anterior (N=3) or posterior (N=3) VMHvl of PR-Cre female mice following the abovementioned stereotaxic coordinates (60nL at a 1:1 ratio). Then, after 14-18 days of AAV expression, Avian Sarcoma Leukosis Virus Subtype A (EnvA) G-deleted rabies virus particles (ΔG-RV) expressing GFP, was injected (100nL) in the same coordinates and the brain was processed, imaged, and analyzed after seven days of ΔG-RV expression.

### Image acquisition and analysis

Images were obtained using an automatic slide-scanning fluorescence microscope (Zeiss Axoscan 7) and were individually exported to tiff format for posterior analysis using the Zen Lite Zeiss software. For all tracing experiments, we imaged the native fluorescence of FG and virally expressed mCherry and GFP without any further amplification. FG, GFP, and mCherry fluorescence were acquired using 405 nm, 488 nm, and 561 nm excitation channels, respectively. For FG tracing experiments, we used the QUINT analysis pipeline^20^, which consisted of four consecutive steps:

(1) Background correction of FG fluorescence was performed using a scaled subtraction approach based on a control channel. Given that all brains in this experiment were scanned using 405 nm and 488 nm excitation channels, for each FG image (I_FG_), we obtained a corresponding “GFP” channel image that contained no specific signal in this experiment to estimate background fluorescence (I_BG_). A scaling factor was calculated as the ratio of the mean pixel intensity of the FG image to the mean pixel intensity of I_BG_. I_BG_ was multiplied by this scaling factor and subtracted from I_FG_ on a pixel-by-pixel basis, yielding a background-corrected FG image (Fig. S1):

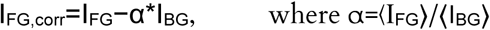

All corrected images were then resized using the Resize function of Nutil to meet the QuickNII input requirement of a maximum image size of 16 MP. The same resize factor was applied to all images.

(2) Image registration to a reference atlas using QuickNII: For each animal, an extensible markup language (XML) descriptor file containing all its brain sections was generated using the “FileBuider.bat” tool provided with QuickNII (RRID:SCR_016865).

(3) Segmentation of FG^+^ cells was performed using Ilastik^21^, which allowed us to train a random forest classifier to distinguish cell objects from the fluorescent signal.

(4) Quantification of features per atlas region was performed by using the Quantifier feature of Nutil^20^. This step generated a set of report files and customized atlas images superimposed with the object pixels (signal).

The output of the QUINT pipeline was analyzed using MATLAB and RStudio. Several exclusion criteria were applied to the dataset: (1) Objects detected with an area of 1 pixel were considered noise and excluded from further analysis. Regions containing 0 object pixels across all animals in the batch were also excluded; (2) Brain areas with subregions were clustered together, summing the object pixels and the area (e.g. ‘Primary motor area, Layer 2/3’, ‘Primary motor area, Layer 5’, ‘Primary motor area, Layer 6a’, ‘Primary motor area, Layer 6b’ were clustered to ‘Primary motor area’). (3) After this, brain regions that had less than 4 detected objects in 2 or more animals (for either anterior or posterior injection batches) were excluded. (4) Signal on fiber tracts was not included in the analysis. (5) The tuberal hypothalamus was excluded from the analysis for being too close to the injection site.

For each brain region we obtained an area (in μm^2^) and the signal (number of objects, FG^+^ cells). With these parameters, we reported two metrics to characterize the afferent connectivity of the VMHvl: First, as employed in previous studies^17^, we describe projections as the percentage of the total input (dividing the number of FG^+^ cells in each area by the total number of FG^+^ cells detected) (Fig. 1B-C, Fig. 4B-C). Second, we report projection density by dividing the number of FG^+^ cells detected in each brain region by its area (projecting cells/μm^2^).

The same QUINT pipeline and exclusion criteria were applied to the raw images containing GFP-labeled neurons from the rabies tracing experiments. Starter cells were manually scored at the AP level corresponding to the injection center. Manual scoring was performed by overlaying the corresponding Paxinos atlas level onto histological images in Adobe Illustrator.

### Statistical analysis

Statistics were performed using RStudio and graphpad software. For comparisons between groups in Figures 1 and 4, we performed a permutation based independence test using the independence_test() function from the RStudio coin package. A random seed of 123 was set to ensure reproducibility, and a Monte Carlo resampling with 10,000 iterations was performed to approximate the p-values. In Figure 3, Two-way ANOVA tests were performed to compare FG labeling density along different brain subnuclei after injections in different AP subdomains of the VMHvl (aVMHvl vs. pVMHvl). We used the Sidak test to correct for multiple comparisons. Statistical significance was set at p < 0.05.

## Abbreviations

AHN: Anterior hypothalamic nucleus
MeA: Medial Amygdala
Rch: Retrochiasmatic area
Arc: Arcuate hypothalamic nucleus
PA/AHIA: Posterior amygdala / amygdalo-hippocampal transition area
LH: Lateral hypothalamus
DMH: Dorsomedial hypothalamus
BMA: Basomedial amygdala
PBN: Parabrachial nucleus
Pe: Periventricular hypothalamus
PP/PIL: Peripeduncular thalamus/posterior intralaminar thalamic nucleus
BNST: Bed nuclei of stria terminalis
POA: Preoptic area
LS: Lateral septum
vHIP: Ventral hippocampus
SPVZ: Subparaventricular zone
PMV: Premammillary ventral
PVH: Paraventricular hypothalamic nucleus
PVA: Paraventricular nucleus of the thalamus
PH: Posterior hypothalamus
PVT: Paraventrincular nucleus of the hypothalamus
AVPV: Anteroventral periventricular hypothalamic nucleus
PMD: Premammillary dorsal
Re: Nucleus of Reuniens
ZI: Zona incerta
PNR: Pontine reticular nucleus
PPN: Pedunculopontine nucleus
PAG: Periaqueductal gray
Str: Striatum
SUM/MM: Supramammillary/mammilary nuclei,
SON: Supraoptic nucleus.

## Acknowledgements

We thank the Champalimaud Foundation (CF) Advanced Bio-imaging and Bio-optics Experimental platform (ABBE) for technical support. This work was supported by the CF through Fundação para a Ciência e a Tecnologia (FCT) Portuguese national funds in the context of the project UIDB/04443/ 2020; by the research infrastructure CONGENTO, co-financed by Lisboa Regional Operational Programme (Lisboa2020), under the PORTUGAL 2020 Partnership Agreement, through the European Regional Development Fund (ERDF) and by the Ramon y Cajal investigator program (RYC2023-043390-I, N.G.C) and the *Proyectos de Generación de Conocimiento 2024* (PID2024-156368NA-I00, N.G.C) grant of the Spanish national funds.

## Author contributions

PRC and JR performed all analyses and made the figures of the manuscript. NGC and DFM performed the tracing experiments and imaging of the samples. LF contributed to the imaging and histological procedures. PD and ICD contributed to the analysis. SQL and NGC conceptualized the work, gathered the funding and wrote the manuscript.

## Supplementary Figures and Tables

**Figure S1.**
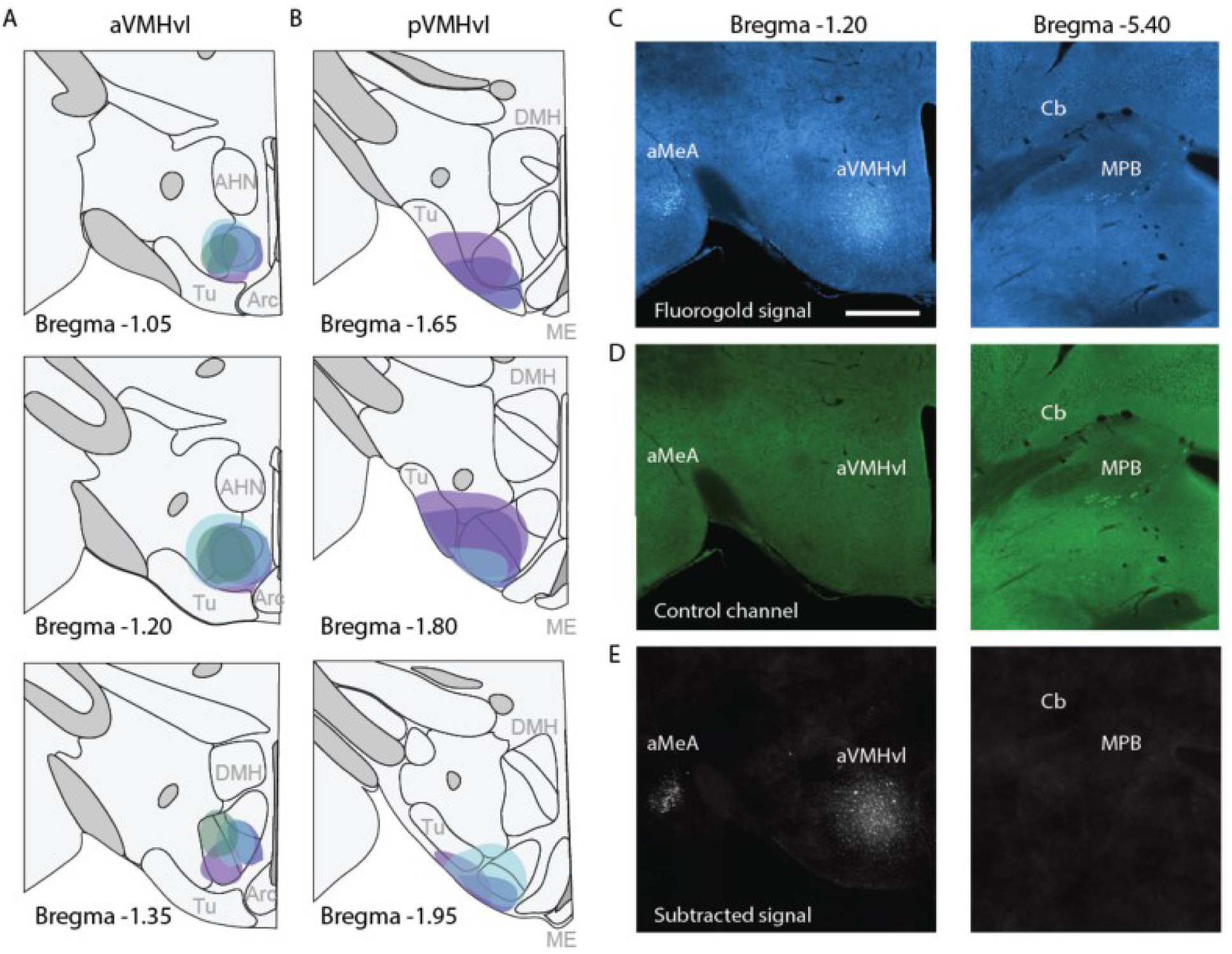
FG injections location and background correction pipeline. FG injections were centered in the aVMHvl (A) or pVMHvl (B) for the different experimental groups. Three AP levels are shown to depict AP injection spread. The raw Fluorogold signal (C) was background-subtracted using a control unlabeled channel (D) to isolate the specific FG fluorescence (E). As an example, the corrected FG signal is visible at the injection site in the aVMHvl and in its transported labeling within the anterior medial amygdala (aMeA) (C-E, left panels). This preprocessing step effectively removed regions in which autofluorescent cells might otherwise be detected by automated pipelines, such as the medial parabrachial nucleus (MPB) (C-E, right panels). The resulting background-corrected images were then processed using the QUINT analysis pipeline for whole-brain detection of FG-positive cells. Scale bar 500 µm.

**Figure S2.**
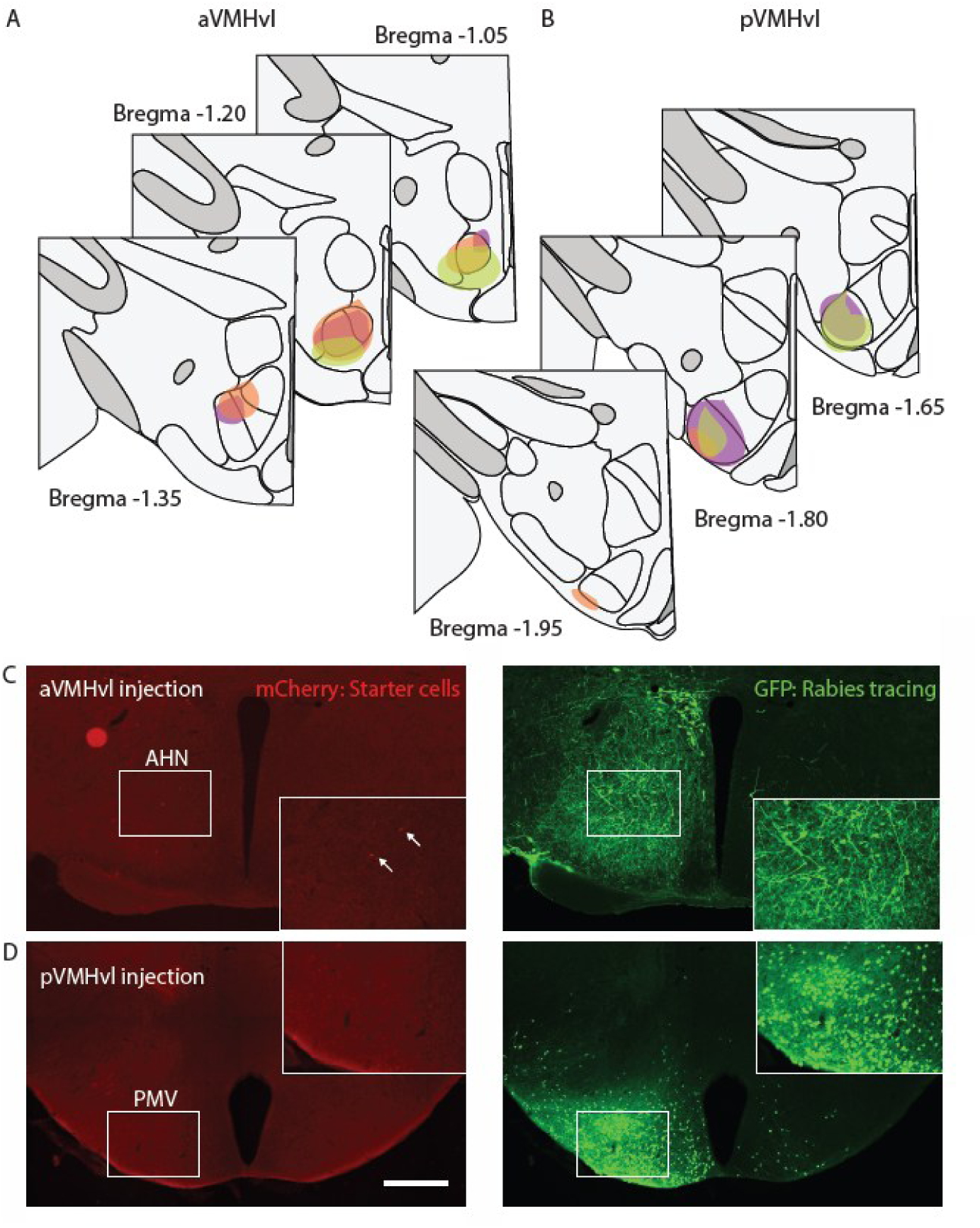
Rabies injection sites and absence of contamination from neighboring regions. Starter virus injections (TVA+mCherry) were centered in the aVMHvl (A) or pVMHvl (B) for the different experimental groups. Three AP levels are shown to depict AP injection spread. Despite their proximity to the injection site, the major sources of input to the aVMHvl and pVMHvl are not attributable to injection bleedthrough. Even in the near absence of starter cells (C-D, left panels), the AHN and PMV remained prominent sources of input to aVMHvl and pVMHvl PR+ neurons, respectively (A-B, right panels). Scale bar 500 µm.

**Figure S3.**
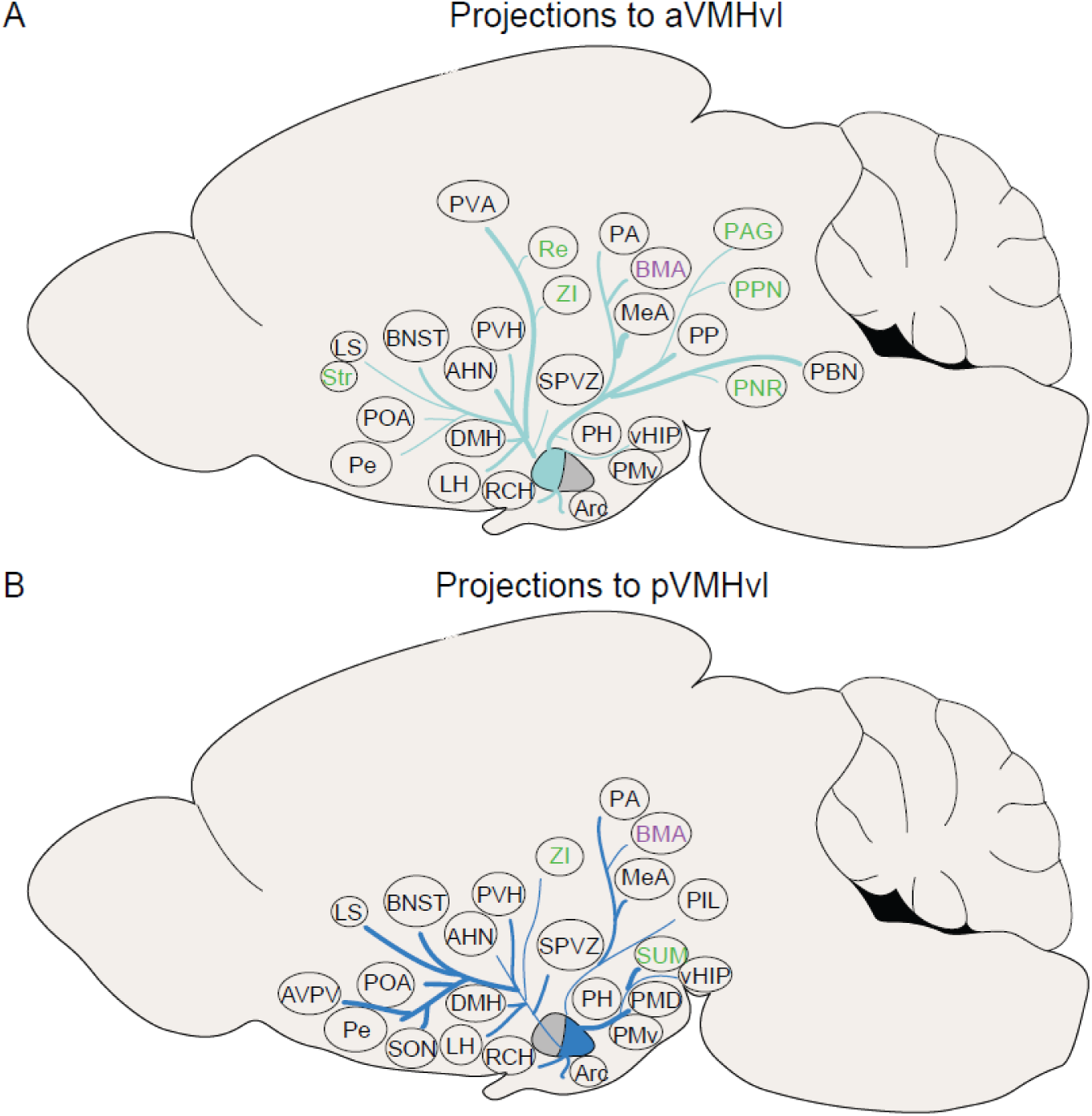
Summary of projections. Semiquantitative schematic summarizing the main input regions to aVMHvl (A) and pVMHvl (B). Arrow thickness indicates relative projection strength. Black labels denote regions consistently identified in both FG and rabies tracing. Green labels indicate regions that did not rank among the top twenty in FG tracing but they did with rabies tracing. Purple labels indicate regions identified in FG tracing but not in rabies tracing.

